# Alterations in excitatory and inhibitory synaptic development within the mesolimbic dopamine pathway in a mouse model of prenatal drug exposure

**DOI:** 10.1101/2020.12.18.423503

**Authors:** Taylor Boggess, James C. Williamson, Ethan B. Niebergall, Hannah Sexton, Anna Mazur, Richard D. Egleton, Lawrence M. Grover, W. Christopher Risher

## Abstract

The rise in rates of opioid abuse in recent years has led to an increase in the incidence of neonatal abstinence syndrome (NAS). Despite having a greater understanding of NAS and its symptoms, there still remains a lack of information surrounding the long-term effects of prenatal exposure to drugs of abuse on neurological development. One potential outcome of prenatal drug exposure that has been increasingly explored is disruption in normal synaptogenesis within the central nervous system. Both opioids and gabapentin, an antiepileptic drug commonly co-abused by opioid abuse disorder patients, have been shown to interfere with the normal functioning of astrocytes, non-neuronal glial cells known to serve many functions, including regulation of synaptic development. The goal of this study was to investigate the effects of prenatal drug exposure on synaptogenesis within brain regions associated with the mesolimbic dopamine pathway, the primary reward pathway within the brain associated with drug abuse and addiction, in a pregnant mouse model. Immunohistochemistry (IHC) and confocal fluorescence microscopy imaging studies on the brains of postnatal day 21 (P21) mouse pups revealed a significant increase in the mean number of excitatory synapses within the anterior cingulate cortex (ACC), nucleus accumbens (NAc), and prefrontal cortex (PFC) in mice that were prenatally exposed to either the opioid drug buprenorphine or gabapentin. These studies also revealed a significant decrease in the mean number of inhibitory synapses within the NAc and PFC of mice treated with buprenorphine. This observed net increase in excitatory signaling capability within the developing mesolimbic dopamine pathway suggests that exposure to drugs of abuse *in utero* can trigger maladaptive neuronal connectivity that persists beyond the earliest stages of life.

## Introduction

The recent rise in rates of opioid abuse in the United States has led to many negative consequences [1]. Increased abuse among pregnant women in particular [2, 3], has led to an increase in incidence of neonatal abstinence syndrome (NAS) [4, 5]. NAS is the clinical diagnosis used to describe the collection of signs and symptoms commonly observed in the newborns of mothers who abused certain drugs, such as opioids, during pregnancy. The developing fetus acquires a physiological dependence on these drugs and, after being separated from the supply of drug at birth, the infant soon displays the symptoms of withdrawal, which can include irritability, tremors, excessive crying, poor feeding, diarrhea, and, in some of the more severe cases, seizures [6]. NAS represents a serious emotional toll for the families of these children and a significant financial burden for healthcare providers [7], prompting considerable research efforts into the pathophysiology of NAS and how to improve the acute treatment of NAS symptoms [6, 8–12]. However, knowledge concerning the long-term effects of prenatal exposure to drugs of abuse on neurological development is limited.

Neurological function is critically dependent on the proper formation of synapses during early development in a process known as synaptogenesis. Various groups have speculated that disruptions in synaptogenesis may underlie many neurodevelopmental and neurodegenerative disorders [13–16]. Potential mechanisms to explain disruptions in synaptogenesis have been proposed, several of which implicate astrocyte dysfunction as a primary factor [17–19].

Astrocytes are the most numerous neuroglia in the central nervous system (CNS) and serve many vital functions, including the ability to regulate the development of synapses within the CNS [20, 21]. Astrocytes promote synaptogenesis in developing neurons by secreting a variety of different factors, including a class of extracellular matrix glycoproteins known as thrombospondins (TSPs) [22]. TSPs (particularly TSP-1 and 2 in developing mammalian brains) exert their synaptogenic effects by binding to α2δ-1, a subunit of L-type voltage gated Ca^+2^ channels, on the surface of neurons at the site of synaptic terminals [13, 22, 23]. Opioid treatment in rodents has been shown not only to decrease astrocyte expression of TSPs [24, 25], but also to interfere with normal astrocyte growth and development [26], indicating that astrocyte-induced synaptogenesis may be key to understanding the effects of opioids on neurological development.

While the increased prevalence of opioid abuse is often credited with the rise in incidence of NAS, it is important to note that many mothers of NAS patients abuse multiple drugs besides, or in combination with, opioids. One prescription drug that has been shown to be increasingly co-abused along with opioids is gabapentin, an anticonvulsant drug also prescribed for the treatment of neuropathic pain [27]. Gabapentin was initially believed to have no potential for abuse or addiction [27, 28]. However, surveys of opioid use disorder patients have found that as many as 26% of those interviewed reported abusing gabapentin for nonmedical reasons, often abusing gabapentin in combination with opioids as a means to potentiate the experienced high [27, 29, 30]. In addition, clinicians have observed a unique presentation of NAS in infants whose mothers abused both opioids and gabapentin while pregnant, with symptoms including tongue thrusting, back arching, and increased eye wandering [31].

The mechanism of action of gabapentin may offer some insight into not only its potential interaction with opioids, but also the unique presentation of NAS associated with the co-abuse of both drugs. Though originally designed as a structural analogue for the neurotransmitter GABA, gabapentin does not bind to GABA _A_ or GABA _B_ receptors. It is instead proposed that gabapentin decreases neurotransmitter release from the presynaptic terminal by inhibiting Ca^+2^ influx through L-type channels [32, 33]. Gabapentin acts to inhibit this Ca^+2^ influx by binding to α2δ-1 [34, 35], the same subunit/receptor through which TSPs mediate astrocyte-induced synaptogenesis. Intriguingly, gabapentin has also been shown to disrupt normal synaptogenesis by interfering with TSP-α2δ-1 binding [23], which leads to the question of whether this signaling is detrimental to CNS development in cases of prenatal gabapentin exposure.

Although the mechanisms through which gabapentin and opioids may interact are unknown, the addictive nature of such drugs of abuse can be for the most part be attributed to their activity within the mesolimbic dopamine pathway (see Figure 1) [36–38]. Examining the effects of drugs of abuse (specifically opioids and gabapentin) on synaptogenesis within brain regions associated with this pathway, specifically the prefrontal cortex (PFC), anterior cingulate cortex (ACC), and nucleus accumbens (NAc), may lead to insight on how these drugs affect neurological development in prenatally exposed children. The hypothesis of this study was that prenatal exposure to buprenorphine, gabapentin, or a combination of both drugs would lead to significant alterations in excitatory and inhibitory synaptic formation in the brain regions of mice associated with addiction and reward.

**Figure 1:**
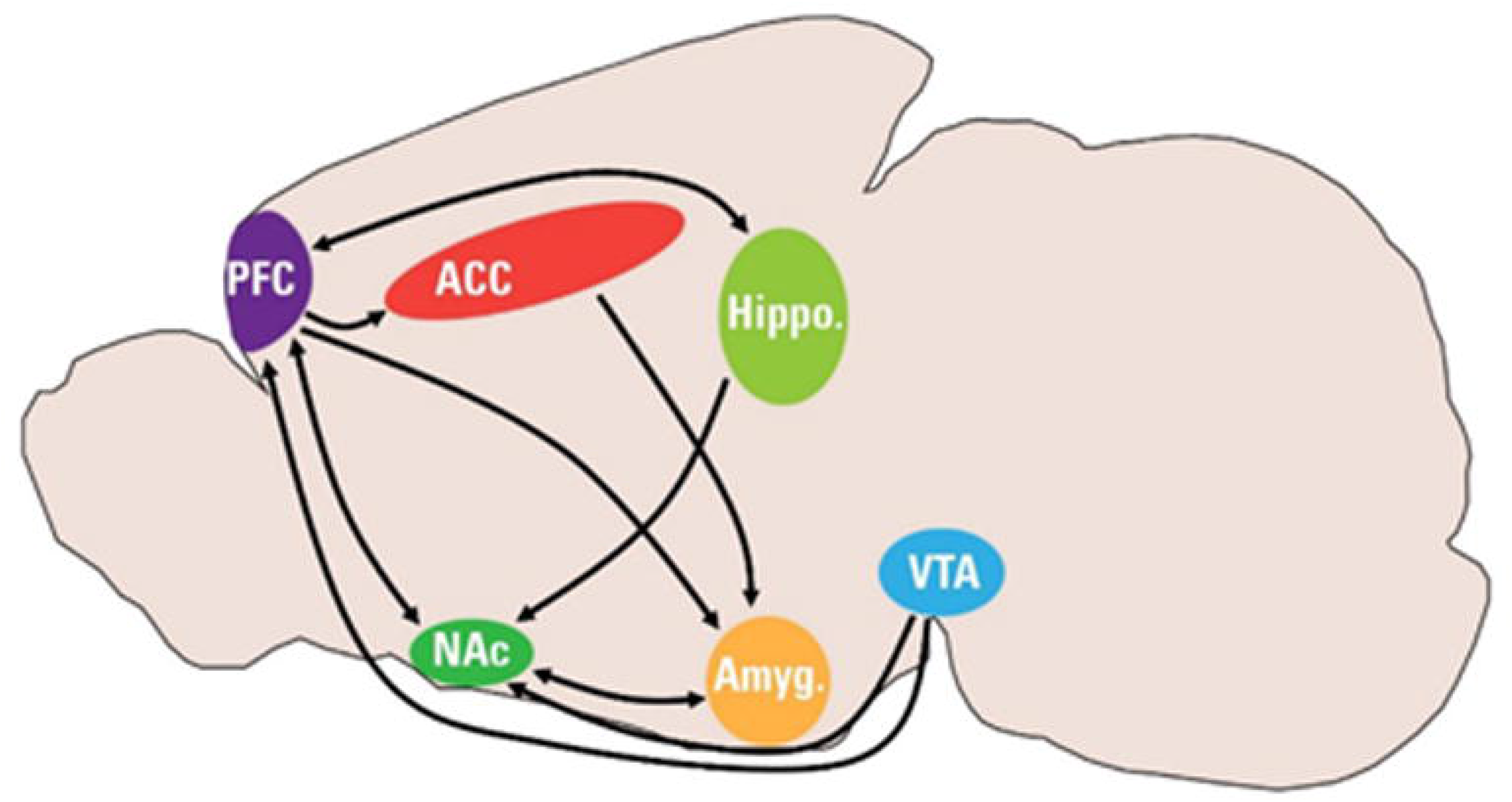
Diagram of mouse brain regions associated with mesolimbic dopamine pathway (PFC: prefrontal cortex, ACC: anterior cingulate cortex, NAc: nucleus accumbens, VTA: ventral tegmental area, Hippo: hippocampus, Amyg: amygdala)

## Methods

### Animals and Drug Treatment

All experiments were conducted in accordance with Marshall University’s Institutional Animal Care and Use Committee (IACUC) guidelines (W.C.R. protocols 696 and 697). Adult C57/Bl6 mice heterozygous for *Cacna2d1* (i.e. α2δ-1 +/−) [13] were a kind gift from Dr. Cagla Eroglu (Duke University). The mice were bred onsite at the Marshall University Animal Resource Facility. Virgin α2δ-1 +/− females and α2δ-1 +/− males were set up as mating pairs. Upon visual confirmation of the vaginal plug (established as embryonic day 0 [E0]), males and females were separated. On E6, pregnant females were given access to 1ml of a 1:1 sweetened condensed milk/ water solution served in a plastic dish. Starting on E7, pregnant females were given free access to a once daily 1ml solution of 1:1 condensed milk/water containing either pharmaceutical grade buprenorphine hydrochloride (CIII) (5 mg/kg; Spectrum Chemical, Gardena, CA), gabapentin (30 mg/kg; Spectrum), a combination of both drug doses, or vehicle control. Drug doses were calculated based on the weight of pregnant females on E7. Daily dosing continued through the birth of pups and ended on postnatal day 11 (P11) (n=11 vehicle control, n=7 30 mg/kg gabapentin [GBP], n=4 5 mg/kg buprenorphine + 30 mg/kg gabapentin [Bup+GBP], n=6 5 mg/kg buprenorphine [Bup]). All animals were given ad libitum food (standard mouse chow milled on site) and water.

### Genotyping

At P7, the tails of the pups were clipped and collected. Toe pads were tattooed (AIMS NEO9 Animal Tattoo System) for later identification. Tissue digestion was performed on the clipped tails and DNA was isolated for PCR amplification and genotyping with a 2% agarose gel. Pups were identified as either α2δ-1 +/−, α2δ-1 +/+ wildtype (WT), or α2δ-1 −/− knockout (KO) using the following primers: F1 (WT forward, 5′-TCTCAGTTACAAGACTATGTGG-3′), F3 (KO forward, 5′-GGCTGTGTCCTTATTTATGG-3′), and LAF-Test (reverse, 5′-AGTAGGAGAAGGTACAATCGGC-3′) (Integrated DNA Technologies, Coralville, Iowa).

### Perfusion, Freezing, and Cryosectioning

At P21, the dam and pups were anesthetized with avertin and cardiac perfusion was performed, first with 0.24μg/ml heparin salt in 0.1M phosphate buffered saline (PBS) for 2 minutes followed by 4% paraformaldehyde (PFA) for 2-5 minutes at a flow rate of 5ml/min. The brains of the dam and pups were extracted and submerged in 4% PFA at 4°C for 24 hours. After 24 hours, the brains were rinsed in PBS and then placed in 30% sucrose:PBS solution for 48 hours. Brains in sucrose solution were frozen within plastic embedding molds (Electron Microscopy Sciences, Hatfield, PA Cat#70182) in 2:1 sucrose solution:Tissue Freezing Medium (Electron Microscopy Sciences, Hatfield, PA Cat#72592-G) and stored at −80°C. Brains were then cryosectioned using a Leica CM 1950 (Leica, Wetzlar, Germany) to 20μm slices and stored in 50% glycerol:Tris-Buffered Saline (TBS) at −20°C.

### Synaptic Labeling Immunohistochemistry

Three independent coronal sections per each mouse containing either the PFC (bregma, 2.96 to 2.58 mm; interaural, 6.76 to 6.38 mm) or the ACC and NAc core/shell (bregma, 1.42 to 0.86 mm; interaural, 5.22 to 4.66 mm) were used for analyses [39]. After selection, brain slices were washed in TBS + 0.2% Triton X-10% (Roche Diagnostics, Mannheim, Germany Cat#11332481001) w/v (TBST) solution at room temperature. The slices were then placed in a 5% normal goat serum (NGS; Jackson Immuno Research Laboratories Inc., West Grove, PA Cat#005-000-121):TBST solution for 1 hour to block nonspecific binding sites. After blocking was completed, the slices were then placed in 5% NGS : TBST containing primary antibodies against both pre- and postsynaptic proteins. In order to label excitatory glutamatergic synapses, anti-vesicular glutamate transporter 1 (VGluT1) antibodies (EMD Millipore, Burlington, MA Cat# AB 5905 guinea pig anti-VGLUT1 polyclonal antibody) were used to bind to presynaptic axonal regions and anti-post synaptic density protein 95 (PSD-95) antibodies (Invitrogen, Carlsbad, CA Cat# 51-6900 rabbit PSD-95 polyclonal antibody) were used to bind to postsynaptic dendritic regions. To label inhibitory GABAergic synapses, anti-vesicular GABA transporter (VGAT) antibodies (Synaptic Systems, Göttingen, Germany Cat# 131004 guinea pig anti-VGAT cytoplasmic domain antiserum) were used to bind to presynaptic axonal regions and anti-gephyrin antibodies (Synaptic Systems Cat# 147002 rabbit anti-Gephyrin polyclonal antiserum) were used to bind to postsynaptic dendritic regions. Slices were incubated overnight on an orbital shaker at 4°C.

After overnight incubation was completed, brain slices were removed from primary antibody solution and washed with TBST. Slices were then placed in 5% NGS:TBST containing fluorescent secondary antibodies against the primary antibodies. Invitrogen Alexa Fluor IgG (H+L) 594 Goat Anti-Guinea pig (Cat# A11076) was used to stain against presynaptic antibodies (either VGluT1 for excitatory or VGAT for inhibitory) and Invitrogen Alexa Fluor IgG (H+L) 488 Goat Anti-Rabbit (Cat# A11034) was used to stain against postsynaptic antibodies (either PSD-95 for excitatory or Gephyrin for inhibitory). Slices were incubated for 2 hours at room temperature in darkness. After incubation was complete, the slices were washed with TBST.

After the final washing step, the slices were rinsed once in 2/3 TBS : 1/3 dH_2_0 solution. Then, the slices were transferred onto glass slides and excess liquid was suctioned off of the slides with a pipet. 1 drop of mounting media containing DAPI (VectaShield, Burlingame, CA Cat#H-1200) was applied to each slice before a cover slip was placed and sealed with clear nail polish. The glass slides were allowed to dry overnight in darkness before being stored at −20°C.

### Confocal Fluorescence Microscopy for Synaptic Protein Imaging

Images of the slides were captured using the Leica SP5 Confocal Fluorescent Microscope housed and maintained by the Marshall University Molecular Biological and Imaging Core. Leica LAS AF software was used to capture 5μm z-stacks with 15 steps (0.33μm distance between each step) within the brain regions of interest across all treatment groups (imaged area/scan = 19,036 μm^2^; 63× oil objective, 1.4 NA).

### Synaptic Puncta Image Analysis

The saved images were analyzed using ImageJ (NIH) software. The custom plug-in “ProjectZ_Triple” (available by request) was used to average and convert the 15 separate images of the z-stack into 5 separate maximum projections, each representing a 1 μm “mini-stack”. Puncta Analyzer (Dr. Cagla Eroglu, Duke University) was then used to quantify the number of discrete puncta for the 488 wavelength channel (corresponding to PSD95 in brain slices stained for excitatory synapses or Gephyrin for inhibitory synapses), the 594 wavelength channel (corresponding to VGluT1 in brain slices stained for excitatory synapses or VGAT for inhibitory synapses), and the colocalized puncta, where puncta from both channels overlapped within 4 pixels of each other. These regions of colocalization of pre- and postsynaptic markers were designated as synapses [40]. The counted synapses were compiled into spreadsheets (Microsoft Excel, Redmond, WA) and analyzed with GraphPad Prism (San Diego, CA).

### Statistical Analysis

Statistical differences were analyzed using one-way ANOVA to detect differences in mean number of synapses. Post-hoc Tukey’s multiple comparisons tests were performed to detect significant differences (p<0.05) between the mean values of each treatment group. Analyses were performed using GraphPad Prism. All values are given as mean +/− standard error of the mean (SEM).

## Results

### Increased Glutamatergic Synapses Following Early Life Drug Exposure

The mean number of neurons counted within the brain regions of interest of all the mouse pups, both male and female, subjected to the same drug treatment was calculated and compared to demonstrate any significant differences. The three brain regions chosen are known to be associated with the mesolimbic dopamine pathway (Figure 1). Also known as the dopamine reward pathway, this pathway works to encode experiences as pleasurable and reinforce future behaviors in an attempt to seek out those rewarding experiences. Opioids are known to interact with this reward pathway by binding to opioid receptors on the surfaces of interneurons within the ventral tegmental area (VTA), a central location for the cell bodies of a large number of dopaminergic neurons. This opioid-receptor interaction inhibits the release of GABA within the VTA, thus disinhibiting dopamine release by dopaminergic neurons [41]. The axons of VTA dopaminergic neurons project to the nucleus accumbens (NAc), where increased phasic release of dopamine in response to external stimuli has been shown to reinforce pleasurable behaviors including eating food, engaging in sexual activity, or taking drugs of abuse [42]. The anterior cingulate cortex (ACC) is involved in a variety of functions, including reward anticipation and modulation of goal oriented motor activity [43, 44], and has been shown to influence motivation via interaction with the mesolimbic dopamine pathway [45]. The prefrontal cortex (PFC) is highly associated with decision making and impulse control and has been shown to have axonal projections to the NAc capable of modulating neuronal activity within this region [46].

First examining excitatory glutamatergic synapses via IHC and confocal microscopy (Figure 2), we observed that P21 mice that were prenatally exposed to each of the three drug treatments showed a significantly higher mean number of VGluT1/PSD95-positive co-localized synaptic puncta than mice treated with a vehicle control (Vehicle control: 284.0 ± 14.21, GBP: 592.0 ± 39.05, BupGBP: 637.6 ± 60.67, Bup: 550.1 ± 37.27). This same trend was also observed within the NAc (Vehicle control: 288.2 ± 21.11, GBP: 631.6 ± 41.54, BupGBP: 627.0 ± 61.35, Bup: 529.2 ±) and the PFC (Vehicle control: 351.9, ± 21.28, GBP: 786.6 ± 44.14, BupGBP: 577.1 ± 66.19, Bup: 923.9 ± 57.89). However, unique to the PFC was a significantly lower mean synapse number in the group treated with both buprenorphine and gabapentin compared to the mouse groups treated with either single drug.

**Figure 2:**
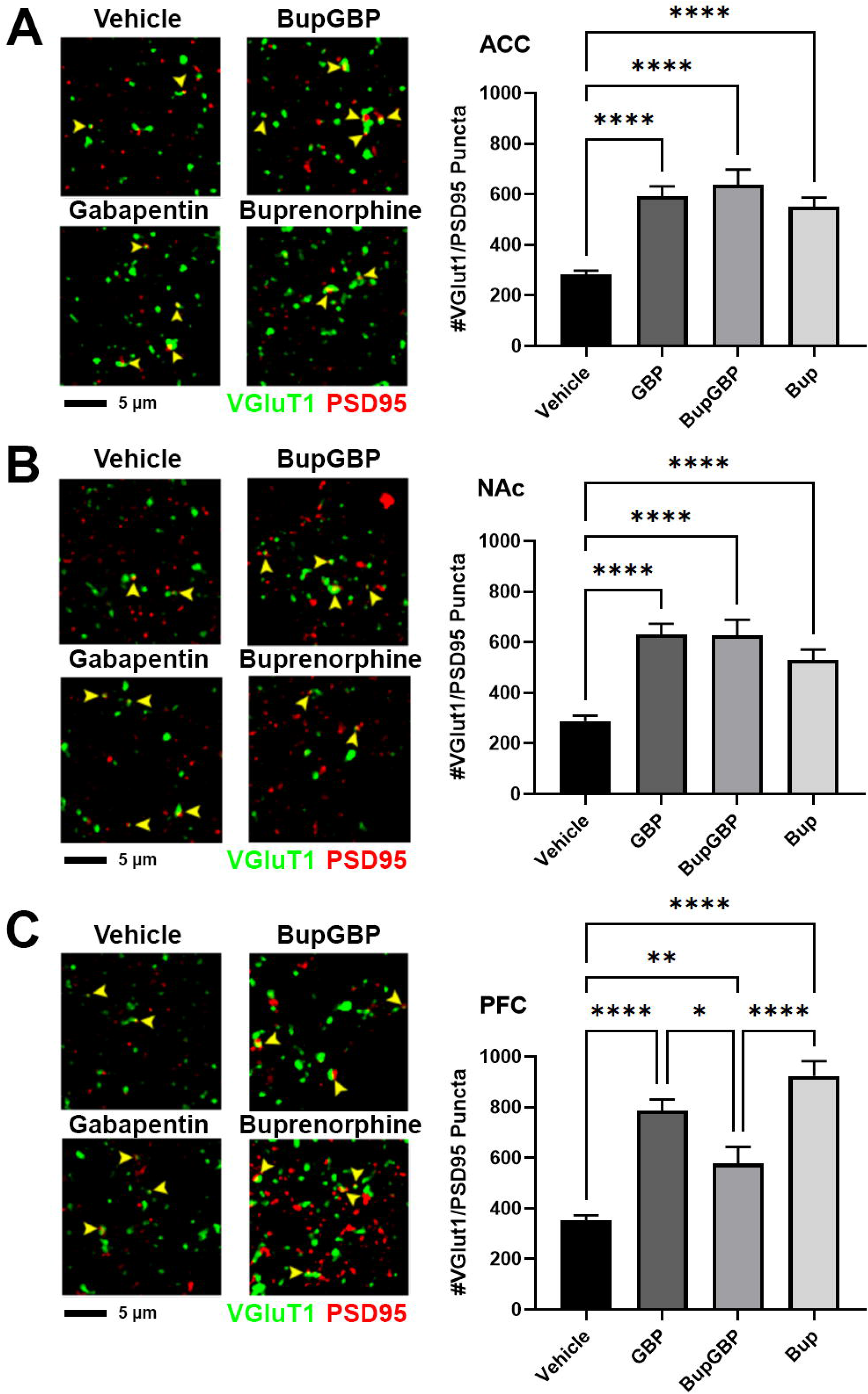
Representative IHC images (left) and quantification (right) of co-localized VGluT1 (green) and PSD95 (red) excitatory synaptic puncta (yellow arrowheads) from prenatal drug-exposed α2δ-1 +/− C57Bl/6J P21 mouse brain within **A)** the anterior cingulate cortex (ACC), **B)** nucleus accumbens (NAc), and **C)** prefrontal cortex (PFC). One-way ANOVA with Tukey’s multiple comparisons post hoc test (ACC: *F*(_3,366_)=21.34; NAc: *F*(_3,366_)=18.81; PFC: *F*(_3,411_)=40.47). n=11 (vehicle control), 7 (30mg/kg gabapentin [GBP]), 4 (5mg/kg buprenorphine + 30mg/kg gabapentin [Bup+GBP]), 6 (5mg/kg buprenorphine [Bup]); *p<0.05; **p<0.01; ****p<0.0001

### Buprenorphine-Induced Decrease in GABAergic Synaptic Development

The same IHC procedure for glutamatergic excitatory synapses was used to stain for GABAergic inhibitory synapses within the same brain regions of the same pups (Figure 3). Within the ACC, no significant differences in mean number of inhibitory synapses were observed between the different treatment groups (Vehicle control: 111.6 ± 10.44, GBP: 112.2 ± 10.86, BupGBP: 127.5 ± 16.93, Bup: 132.5 ± 9.164). However, within the NAc, a significant decrease in inhibitory synapses compared to the vehicle control group (120.8 ± 9.60) was observed in the pups treated with both drugs (63.03 ± 8.10) and in the pups treated with buprenorphine alone (46.18 ± 4.20). Synapse number in the group treated with both drugs was also significantly lower than in the group treated with gabapentin (115.0 ± 7.19) alone. Within the PFC, the only significant difference observed was a significant decrease in synapse number in the group treated with buprenorphine (88.75 ± 6.45) compared to the vehicle control group (209.9 ± 17.57). No significant differences were observed between the vehicle control group and either the group treated with both drugs (153.6 ± 14.97) or the group treated with gabapentin (211.4 ± 13.09). Taken together, these results indicate a general increase in excitatory synaptic signaling within all three brain regions as a result of prenatal exposure to buprenorphine and/or gabapentin.

**Figure 3:**
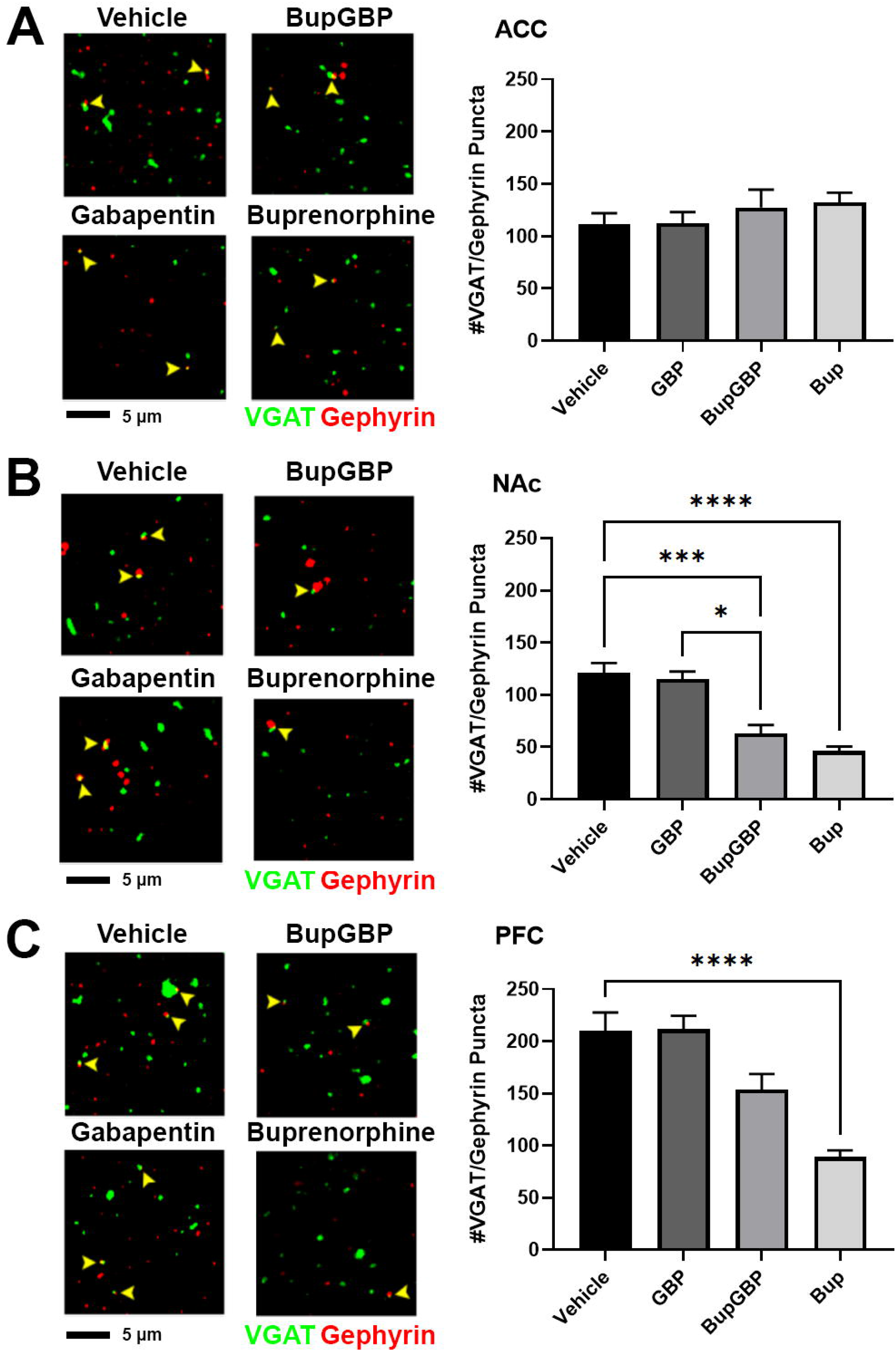
Representative IHC images (left) and quantification (right) of co-localized VGAT (green) and gephyrin (red) inhibitory synaptic puncta (yellow arrowheads) from prenatal drug-exposed α2δ-1 +/− C57Bl/6J P21 mouse brain within **A)** ACC, **B)** NAc, and **C)** PFC. One-way ANOVA with Tukey’s multiple comparisons post hoc test (ACC: *F*(_3,332_)=0.5893; NAc: *F*(_3,336_)=12.26; PFC: *F*(_3,341_)=8.590). n=11 (vehicle control), 7 (GBP), 4 (Bup+GBP), 6 (Bup); *p<0.05; **p<0.01; ****p<0.0001

## Discussion

The goal of this study was to demonstrate the effects of prenatal drug exposure on synaptic development within the mesolimbic dopamine pathway in the mouse CNS. All three drug treatments (buprenorphine, gabapentin, buprenorphine + gabapentin) produced significant increases in mean excitatory synapse number within each of the three brain regions. Regarding inhibitory synapses, the buprenorphine treatment groups showed significant decreases in mean synapse number within the NAc and PFC compared to the vehicle control groups. The double-drug treatment group also showed a significant decrease within the NAc. Finally, quantification of NeuN-expressing cells in these regions revealed that the observed changes in synapse number were not merely due to changes in the number of neurons.

Initially, the observation that prenatal exposure to gabapentin produced an increase in excitatory synapses would appear to contradict the findings of previous studies, including our own, which showed that gabapentin inhibits normal excitatory synapse development by interfering with normal TSP-α2δ-1 binding [23]. One possible explanation for our current finding is that the observed increase in synapse number may be compensatory for an overall decrease in glutamate release. Glutamate signaling is critical for optimal CNS development, while one of the known mechanisms of action of gabapentin is to inhibit neurotransmitter release. Prenatal stress has also been shown to alter expression of glutamate receptors within the brains of rats [47]. Therefore, it is possible that more excitatory glutamatergic synapses are formed within the CNS in order to overcome the lack of glutamatergic signaling during this relatively plastic developmental period. This mechanism may also potentially explain a surprising finding from one of our previous studies, wherein neurons that lacked α2δ-1in presynaptic terminals (partially mimicking the action of gabapentin) actually formed more glutamatergic synapses than those where α2δ-1 signaling was intact [13].

The drug treatments used in this study were chosen not only to produce significant results in the animal subjects, but also to attempt to replicate both the drug doses and timeframes of abuse that would be possible in human mothers. Buprenorphine was chosen for this study due to its increased use in medication assisted therapy (MAT). Pregnant mothers who are addicted to opioids commonly undergo MAT which combines less-addictive prescription opioid drugs— namely methadone or buprenorphine—with behavioral counseling in order to treat addiction and achieve better health outcomes for the mother and child. However, emerging evidence asserts that any opioid use during pregnancy, even when part of a medical treatment plan, can have deleterious effects on the developing fetus [48, 49]. Since all opioid drugs have similar mechanisms of action, they all have the potential to affect the developing CNS. Martin et al. showed that 5mg/kg buprenorphine was necessary to produce significant increases in pain threshold in rats [50]. The dosing for gabapentin in this study was chosen after Kilic et al. found 30mg/kg gabapentin was capable of producing analgesia in mice [51]. The results of this study show that the chosen doses of buprenorphine and gabapentin were sufficient to produce significant changes in synaptic structure within the brains of treated mice.

The dosing schedule, starting on E7 and ending on P11, was chosen to correspond to a period of significant synaptogenesis within mice [22, 52, 53], a period which also roughly correlates to the second trimester of human fetal development [54, 55]. At birth, a mouse pup is considered to be at a developmental stage similar to that of a human fetus in the late second trimester. Despite the fact that newborn mouse pups are no longer receiving direct exposure to drugs of abuse via the placental blood supply, there is still reason to believe that the brains of those pups are still being exposed to those drugs. Kongstorp et al. demonstrated that buprenorphine accumulates and remains in the brain tissue of newborn rodents for several days following birth [56]. In addition, both buprenorphine and gabapentin have been shown to pass from mother to infant via breast milk, albeit in low concentrations compared to the mothers’ blood plasma levels [57–59]. The results of this study show that the chosen treatment schedule is capable of altering normal CNS development in mouse pups.

The pups used in this study were all heterozygous for expression of *Cacna2d1* (i.e. α2δ-1 +/−). Originally, the goal of this study was to mate α2δ-1 +/− C57 Bl6 males and females together to produce pups of all three possible α2δ-1 genotypes: α2δ-1 +/−, α2δ-1 +/+ (wildtype; WT), and α2δ-1 −/− (knockout; KO). There was an initial objective to compare IHC results between the three different genotype groups. It was expected that the absence or presence of α2δ-1 would correlate with significant differences in synapse number, particularly when pups were exposed to the α2δ-1 ligand, gabapentin. However, after several attempts to breed α2δ-1 KO pups, it was found that such pups were either not being born or did not survive to age P21. In fact, in the 6 different litters produced for this study, only one α2δ-1 KO pup (treated with 5 mg/kg buprenorphine) survived to be included in IHC experiments. It was speculated that this low number of viable α2δ-1 KO offspring was due to decreased survivability among these pups. Given the widespread presence of α2δ-1 within the mouse CNS [60, 61], skeletal muscle [62, 63], and cardiac muscle [64], the global knock out of this Ca^2+^ channel subunit in combination with exposure to drugs of abuse during fetal development may have been potentially lethal. This is a potential hurdle that must be cleared in future investigations into the role of α2δ-1 in a model of prenatal drug exposure.

A key finding of this study was the change in the balance of excitation and inhibition present within the analyzed brain regions. Alterations in this balance have been observed in a number of neurological based disorders and developmental disabilities such as Down Syndrome [65], Rett Syndrome [66, 67], and Autism Spectrum Disorders [68–70]. Astrocytes, through a variety of methods, have been shown to influence and maintain this balance [71]. Not only are astrocytes able to mediate both excitatory [22] and inhibitory [72] synaptogenesis, they are also able to mediate synapse elimination via phagocytosis mediated by astrocytic cell surface receptors such as MEGF10 and MERTK [73]. Outside of synapse regulation, astrocytes are also capable of both uptake [74–76] and release [77, 78] of glutamate and GABA, the primary excitatory and inhibitory neurotransmitters within the mammalian brain, respectively. Greater understanding of the role of astrocytes in maintaining the excitatory/inhibitory balance may lead to greater understanding of neurological dysfunction and, when coupled with increased understanding of the effects of drugs of abuse on astrocytes during development, correlations between prenatal drug exposure and neurological dysfunction may be made.

The results of this study indicate a net increase in excitatory signaling capability within the mesolimbic dopamine pathway of mice prenatally exposed to drugs of abuse. Previous studies examining dopamine signaling within the NAc and VTA have shown that phasic dopamine release selectively modulates excitatory but not inhibitory responses of NAc neurons during sucrose-seeking behavior, indicating an influence of dopamine on goal-directed actions [79]. Greater synaptic transmission within the NAc is expected to be indicative of greater excitation in response to a rewarding stimulus. This could indicate a greater level of reinforcement when presented with a rewarding stimulus. Individuals with a greater response to a rewarding stimulus may be at greater risk for developing drug addiction or other disorders resulting from poor impulse control.

In conclusion, this study utilized a mouse model of prenatal drug exposure to examine the effects such exposure may have on synaptogenesis within brain regions associated with reward and addiction. Mouse pups prenatally treated with buprenorphine, gabapentin, or a combination of both drugs were shown to have a greater number of excitatory synapses within these brain regions compared to pups treated with a vehicle control. Pups prenatally treated with buprenorphine were also shown to have a lower number of inhibitory synapses within these same regions. Such changes in synapse number may indicate a general increased in net excitation in response to a rewarding stimulus. These findings may inform future studies examining the relationship between drug addiction and poor impulse control with a history of prenatal drug exposure and NAS.

## Acknowledgements

We would like to thank Mr. David Neff, Dr. Michael Norton and the Molecular and Biological Imaging Center at Marshall University. We would also like to thank Dr. Kelly Hopper and the staff of the Marshall University Animal Resource Facility for excellent veterinary assistance.

This research was funded by the John and Polly Sparks Foundation, the Brain and Behavior Research Foundation NARSAD Young Investigator Award 27662, and to W.C.R. Support was also provided by the Marshall University Genomics and Bioinformatics Core, the West Virginia IDeA Network of Biomedical Research Excellence (WV-INBRE) grant (P20GM103434), the COBRE ACCORD grant (1P20GM121299), and the West Virginia Clinical and Translational Science Institute (WV-CTSI) grant (2U54GM104942).

## Author contributions

T. Boggess, R.D. Egleton, L.M. Grover, and W.C. Risher designed the study. T. Boggess, E.B. Niebergall, H. Sexton, and A. Mazur managed the mouse colony and performed genotyping. T. Boggess, J.C. Williamson, and E.B. Niebergall conducted the experiments. T. Boggess and W.C. Risher analyzed the data and wrote the manuscript.

## Notes

### Competing Interest Statement

The authors have declared no competing interest.

